# Variable sensitivity to DNA damaging chemotherapeutic modulated by cell type-dependent bimodal p53 dynamics

**DOI:** 10.1101/149013

**Authors:** Ruizhen Yang, Bo Huang, Yanting Zhu, Yang Li, Feng Liu, Jue Shi

## Abstract

Mechanisms that determine drug sensitivity of distinct cancer types is poorly understood for most cytotoxic chemotherapy. In this study, we elucidated a new resistance mechanism to DNA damaging chemotherapeutic through modulation of p53 dynamics. While both sensitive and resistant cancer cell lines activated similar p53 oscillation followed by cell-cycle arrest in response to low dose of DNA damaging drug, they switched in a bimodal manner to monotonic or single pulse dynamics at high drug dose. Cell lines with monotonically increasing p53 underwent rapid and extensive drug-induced apoptosis, while those exhibiting a single p53 pulse mostly survived. By combining single cell imaging with computational modeling, we characterized a regulatory module involving ATM, p53, Mdm2 and Wip1, which generates bimodal p53 dynamics through coupled feed-forward and feedback, and we found that basal expression of ATM determined the differential modular output between drug sensitive and resistant lines. Moreover, we showed combinatorial inhibition of Mdm2 and Wip1 was an effective strategy to alter p53 dynamics in resistant cancer cells and sensitize their apoptotic response. Our results point to p53 pulsing as a potentially druggable mechanism that mediates resistance to cytotoxic chemotherapy.

## Introduction

Tumors exhibit large intrinsic variation in drug responsiveness due to both intra-tumoral and inter-tumoral heterogeneity; and previously sensitive tumors commonly evolve to be drug resistant during chemotherapy. In order to improve therapy, we need better understanding of both intrinsic and acquired drug resistance. Most characterized and known mechanisms of drug resistance involve genetic mutation, such as mutation of drug target genes for kinase inhibitors and mutation of genes that mediate drug responses, e.g. p53 for DNA damaging drugs [1, 2]. However, for most commonly used cytotoxic chemotherapy, the mechanistic basis of drug sensitivity is unclear at both genotype and phenotype levels.

The p53 pathway, consisting of the tumor suppressor protein p53, its upstream regulators and downstream target genes, plays a central role in mediating cellular response to DNA damaging chemotherapeutics [3-5]. Loss of p53 function due to p53 mutation has been widely studied with respect to tumor resistance to DNA damaging drugs. However, resistance is often seen, and may be even greater, in tumors with wild-type p53 for unclear reasons [6-8]. Extensive biochemical and cell biology studies have revealed p53 activity is regulated by multiple post-translational modifications [9, 10], differential subcellular localization [11], and interaction with co-factors [12]. Drug resistance may arise from all of these venues. More recently, single cell studies both by us and others showed that alteration of the induction dynamics of p53 is another mechanism to modulate p53 activity, raising the interesting possibility of a mechanistic link between p53 dynamics and drug sensitivity [13, 14].

The control of p53 dynamics in response to DNA damage was first examined for transient gamma or UV irradiation and revealed intriguing oscillation of p53 level that culminated in cell-cycle arrest and senescence, but not cell death [15-18]. Moreover, a recent study of a cancer cell line panel showed damage-induced p53 dynamics varied significantly between cell lines [19]. However, cell-cycle arrest remained the dominant damage response phenotype, despite the difference of p53 dynamics in the distinct cell lines. The lack of cell death response in these previous studies was probably due to the low level of DNA damage used. By examining a much larger dose range of DNA damage triggered by a DNA damaging chemotherapeutic, etoposide, we uncovered a new dynamical regime of p53 activation at high DNA damage, i.e., monotonic induction, which triggers cell death by enhancing p53 activation level and pro-apoptotic activity [13]. We further showed that changing p53 dynamics from oscillation to monotonic increase was sufficient to switch response from cell-cycle arrest to apoptosis in drug sensitive cell lines, confirming the role of bimodal p53 dynamics in regulating differential phenotypic response, i.e., cell-cycle arrest vs. death, to DNA damage. However, it left unexplained the role of p53 dynamics in drug resistance, in particular resistance to drug-induced cell death that is critical for therapeutic efficacy.

We thus set out to compare single-cell p53 dynamics of cancer cell lines that are intrinsically sensitive vs. resistant to DNA damaging drugs. Intriguingly, we observed cell-type dependent bimodal regulation of p53 dynamics that correlates with sensitivity to drug-induced apoptosis. Cell lines sensitive to etoposide, such as U-2 OS and A549, switched from oscillatory p53 to strong monotonic p53 induction followed by rapid cell death, as etoposide concentration increased. In contrast, resistant cell lines, such as MCF7, switched to single p53 pulses at high drug dose, which mostly triggered cell-cycle arrest but not cell death. To unravel the mechanistic basis underlying cell-type variation in p53 dynamics and its role in drug resistance, we combined single cell imaging, ensemble profiling and computational modeling. Our results revealed a multi-component control of coupled feed-forward and feedback involving ATM, p53, Mdm2 and Wip1, which can activate differential p53 dynamics and drug response in a cell-type specific manner. And our data also revealed network perturbations that can revert drug resistance and could be developed into therapeutic strategies.

## Results

### Cell-type variation in p53 dynamics and cellular response

Etoposide is a DNA damaging chemotherapeutic commonly used in the clinic and triggers DNA double-stranded breaks by inhibiting topoisomerase II [20]. We first compared dose response of an etoposide-sensitive (U-2 OS) and resistant cell line (MCF7), and their respective drug-induced p53 dynamics. Both lines have wild-type p53. To monitor real-time p53 dynamics in individual cells, we used clonal fluorescent reporter cell lines that stably express a p53-Venus construct (i.e. wild-type p53 tagged with a yellow fluorescent protein, Venus) [13]. Dynamics of the p53-Venus construct have been confirmed to behave similarly to its wild-type counterpart in both U-2 OS and MCF7 in previous studies [13, 15]. Time-lapse imaging of the fluorescent reporter lines showed that upon drug treatment, the fluorescent signal of p53-Venus mostly localized in the nucleus, and the time course of p53 fluorescence quantified by integrating Venus fluorescence in the nucleus showed a dose-dependent bimodal regulation (Fig. 1a and 1b). At low drug dose (1μM - 10μM), nuclear p53 mostly exhibited periodic pulsing followed by cell-cycle arrest in both cell lines. As etoposide concentration, and the extent of DNA damage, increased, p53 switched from periodic pulsing to different modes that varied between cell lines. High etoposide activated monotonic induction of p53 in U-2 OS cells, with peak level 5-10 fold higher than the amplitude of the pulsing mode (Fig. 1a). In contrast, MCF7 cells switched to a single p53 pulse, with an amplitude 2-3 fold higher than that at low dose and a period about 1.5 fold longer (Fig. 1b). These single-pulse dynamics resulted in a substantially lower integrated induction level of p53 in MCF7 than U-2 OS cells under high drug.

**Figure 1.**
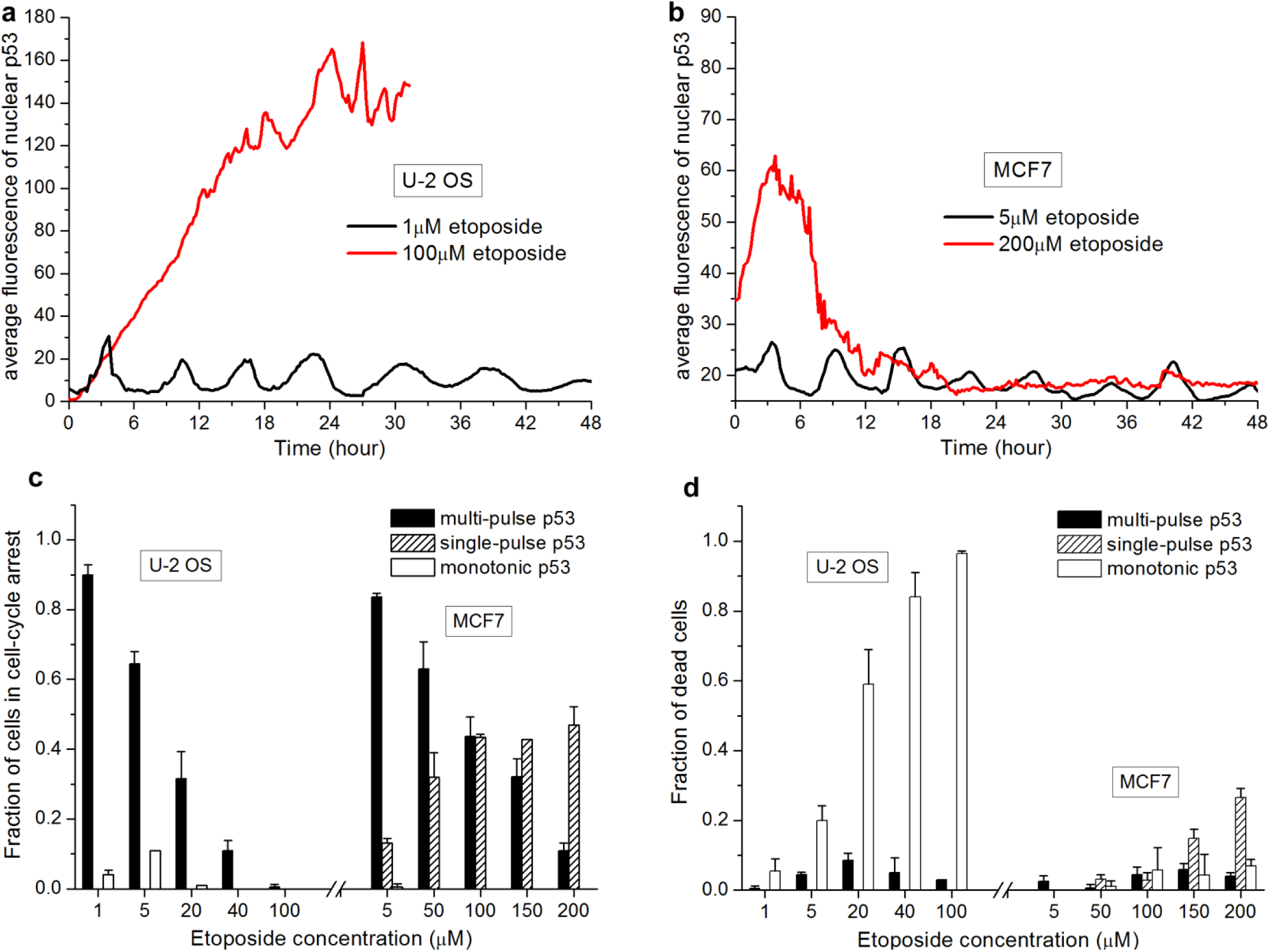
p53 dynamics and cellular response to etoposide are both dose-dependent and cell-type dependent. (a, b) Representative single-cell trajectories of p53 dynamics at the nucleus quantified from fluorescence of the p53-Venus reporter in (a) U-2 OS and (b) MCF7 cells at the indicated low (black lines) and high (red lines) etoposide dose. Cells were treated with etoposide at time 0 and tracked for 72 hrs or till cell death occurred. The abrupt end of p53 trajectories in U-2 OS cell before 72 hr corresponds to the time of death. (c, d) Percentage (±SD) of U-2 OS (left columns) and MCF7 cells (right columns) that exhibited the three distinct p53 dynamics, i.e., multiple pulse (black columns), single pulse (lined columns) and monotonic increase (empty columns) and went into (c) cell-cycle arrest or (cell death) under the indicated etoposide treatment. Data were averaged from two independent sets of single-cell imaging experiments. Total number of cells analyzed ranges from 77 to 113, varied between conditions. Error bars indicate standard deviation. The U-2 OS data were re-plotted from Figure 2b published in reference [13] for direct comparison with the new MCF7 data.

Based on the time-lapse movies, we can score dose-dependent cell fate response by morphological tracking (refer to Materials and Methods for details) and correlated p53 dynamics with the cell fate response in individual cells. We re-plotted dose response data previously published in reference [13] for U-2 OS cells in Figure 1c and 1d for direct comparison with the new data for MCF7 cells. These single cell statistics demonstrated that > 92% MCF7 cells (summed over all drug concentrations) that showed periodic pulsing of p53 went into cell-cycle arrest (Fig. 1c). The strong correlation between oscillatory p53 and cell-cycle arrest is similar to our previous observation in U-2 OS cells, further indicating the role of p53 oscillation in restraining p53 induction level and pro-apoptotic activity so as to render cell-cycle arrest that allows damage repair at low DNA damage. As etoptoside concentration increased, both U-2 OS and MCF7 showed increase of cell death, but MCF7 cells were much less sensitive to etoposide-induced cell death than U-2 OS cells (Fig. 1d). 72-hour treatment of 200μM etoposide only induced about 30% cell death in MCF7, as compared to nearly 100% U-2 OS cell death after only 36-hour treatment of 100μM etoposide, the saturating dose for cell death response for U-2 OS cells. Moreover, in contrast to a strong correlation between monotonic increase of p53 and cell death in U-2 OS cells, cell-cycle arrest remained the dominant cell fate following change of p53 dynamics from multiple pulses to a single pulse in MCF7 at high drug dose.

The observed resistance of MCF7 cell to drug-induced cell death, even in response to very high concentration of etoposide, correlated with the significantly lower p53 induction level rendered by the single-pulse dynamics of p53, as compared to the strong monotonic increase seen in U-2 OS (Fig. 1a & 1b). Among MCF cells that showed single-pulse p53, 80% were found to go into cell-cycle arrest, while the other 20% died. The induction level of single-pulse p53 was obviously not sufficient to cross the threshold for triggering cell death for the majority of MCF7 cells. This observation further indicated the suppressive role of p53 pulsing in attenuating p53 induction level and subsequent cell death. Note that although the MCF7 cells are isogenic, the threshold for triggering cell death may still vary between individual cells, e.g., due to stochasticity in gene expression [21, 22]. Therefore, cell-to-cell variability in sensitivity to cell death coupled with the intermediate level of p53 induction resulted from single-pulse p53 likely account for p53 inducing alternative cell fate of arrest and death in individual cells. Overall, the observation of cell line-dependent regulation of p53 dynamics and its correlation with differential drug response, in particular the cell death response, point to the importance of bimodal p53 dynamics in regulating sensitivity of distinct cancer cells to DNA damaging chemotherapeutics.

### Cell line variation in p53 dynamics at high drug concentration correlates with differential levels of phospho-ATM, Mdm2 and Wip1

To explore the molecular mechanism underlying dose-dependent bimodal regulation of p53 dynamics as well as its cell-type dependence, we profiled, by western blotting, the level of expression and/or post-translational modification of key regulators that are known to modulate p53 expression and activity, including ATM, Chk2, Mdm2 and Wip1 [9, 23-25]. We chose to compare cellular responses at one low (i.e., 1μM etoposide for U-2 OS and 5μM for MCF7) and one high drug concentration (i.e., 100μM etoposide for U-2 OS and 200μM for MCF7) for each cell line, under which cells largely exhibited the same p53 dynamics, i.e. periodic pulsing at low drug and monotonic increase or single pulse at high drug. We performed the western blots at 4 time points after etoposide addition, and ran gels either with the two different drug concentrations side by side for one cell line (Fig. 2a & 2b), or different cell lines side by side for the same dose range (Fig. 2c & 2d), to allow quantitative comparisons.

**Figure 2.**
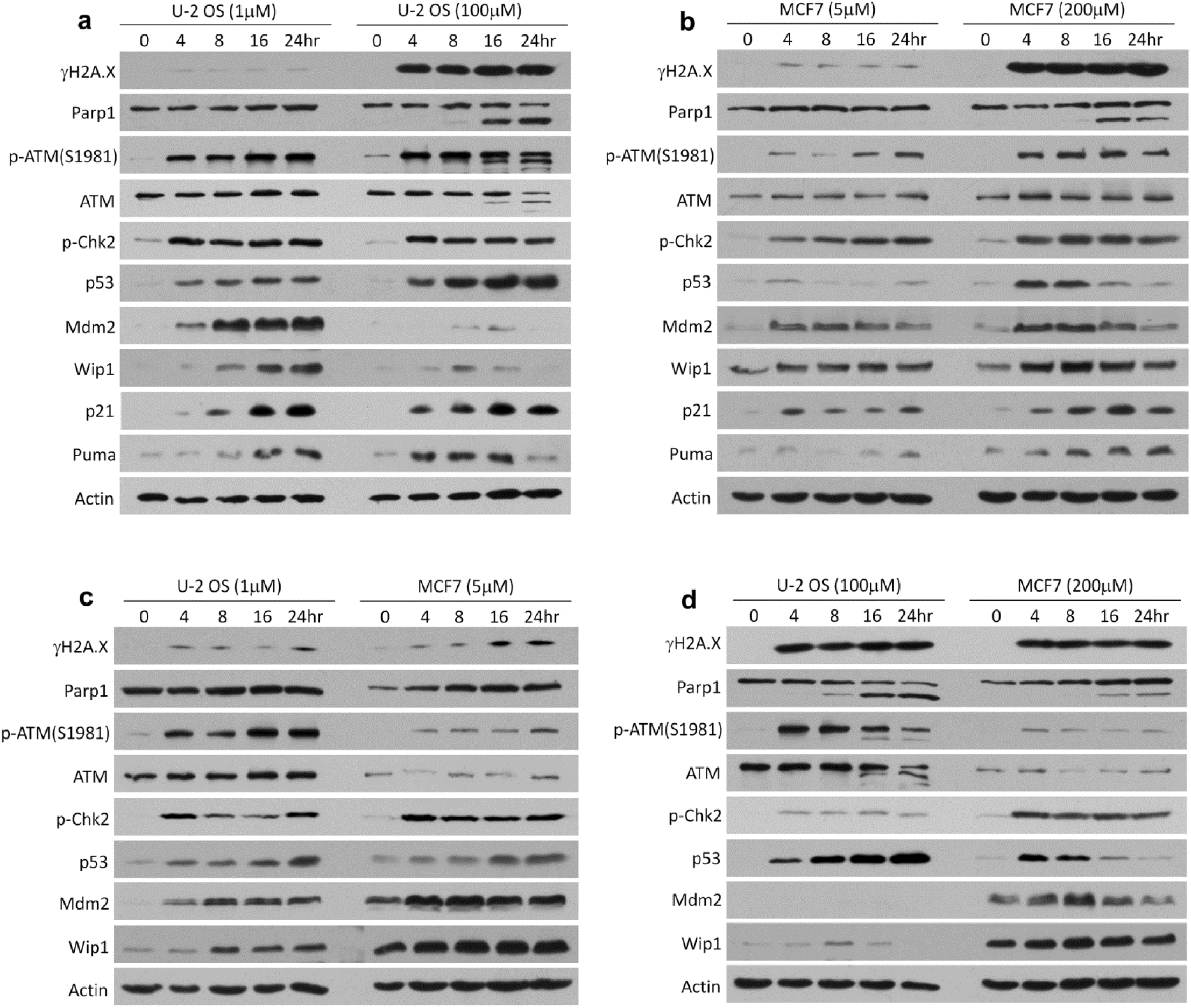
p53 dynamics and p53-mediated drug response correlate with kinetics of a number of proteins/protein modifications. (a, b) Western blot comparison of low vs. high DNA damage in (a) U-2 OS cells and (b) MCF7 cells. (c, d) Comparison of U-2 OS and MCF7 cells at (c) low DNA damage and (d) high DNA damage. For all the western blotting analysis, actin served as the loading control. DNA damage level was indicated by a DNA damage marker, γH2A.X and the extent of cell death was indicated by Parp1 cleavage.

As shown in Figure 2a and 2b, in both U-2 OS and MCF7 cells, higher drug concentration induced higher level of p53, similar to data from single cell imaging (DNA damage level was indicated by γH2A.X, a double-stranded break reporter). And higher p53 level correlated with higher p53 activity in activating its transcriptional targets, such as p21 (cell-cycle arrest gene) and Puma (cell death gene), again indicating bimodal p53 dynamics control cell fate response mainly by modulating p53 induction level. Among the immediate p53 regulators, phosphorylation of ATM at Serine 1981 was more highly induced in response to high drug dose for both cell lines. In contrast, the dose-dependent expression of Mdm2 and Wip1, the two main negative regulators and transcriptional target genes of p53, varied significantly between the two cell lines. For U-2 OS cells, expressions of both Mdm2 and Wip1 steadily increased in time at low etoposide concentration, but such up-regulation was suppressed at high drug (Fig. 2a). The inhibition of negative feedbacks from Mdm2 and Wip1 likely enables p53 to accumulate to a high level, resulting in monotonically increasing p53 in U-2 OS cells. MCF7 cells, however, exhibited strong up-regulation of Mdm2 and Wip1 in response to both low and high drug dose (Fig. 2b). This up-regulation of Mdm2 and Wip1 correlated with the much lower level of p53 induction in MCF7 cells at high drug. Cell line comparison in Figure 2c and 2d further confirmed the differential expression of Mdm2 and Wip1 in U-2 OS and MCF7. It also revealed a much lower level of total ATM and thus lower level of phospho-ATM, but higher basal expression of Wip1, in MCF7 than U-2 OS, which may also contribute to the attenuated p53 induction in the form of single pulse in MCF7 at high drug. We note the level of DNA damage engendered by etoposide in the same dynamical regime, as indicated by the γH2A.X signal, was largely similar in U-2 OS and MCF7 (Fig. 2c and 2d). Therefore, the cell line-dependent dynamics of p53 and its regulators were not due to difference in the input DNA damage signal induced by etoposide, but rather variable activation of the p53 regulatory pathway.

### ATM-mediated Mdm2 degradation accounts for cell line-specific Mdm2 attenuation at high drug concentration

Given the level and dynamics of phospho-ATM, Mdm2 and Wip1 correlated with bimodal p53 dynamics as well as cell-type variation, we next examined their mechanistic involvement in regulating p53 induction. We had previously shown that in U-2 OS cells, attenuation of Mdm2 activity alone, by either gene knockdown or a small-molecule Mdm2 inhibitor, Nutlin-3, was sufficient to change not only p53 dynamics from oscillation to monotonic increase but also cellular response from cell-cycle arrest to cell death [13]. The data thus suggest differential Mdm2 up-regulation modulated by DNA damage level is the major regulator of bimodal p53 dynamics in U-2 OS cells, with Wip1 playing a negligible role in this regard. The fact that U-2 OS cells express significantly less Wip1 than MCF7 cells also pointed to a relatively minor role of Wip1 in U-2 OS, as compared to MCF7.

Suppression of Mdm2 up-regulation in U-2 OS at high drug concentration (i.e., high DNA damage level), which was not seen in MCF7 cells, could be due to reduced gene transcription or translation, or increased protein degradation. We had previously performed quantitative real-time PCR (qPCR) analysis and found no differential transcription of the Mdm2 gene at low vs. high DNA damage in U-2 OS cells [13], indicating post-transcriptional mechanism likely accounts for the dose-dependent Mdm2 expression. DNA-damage kinases, such as ATM, had been observed to promote auto-degradation of Mdm2 [26], which led us to hypothesize that the strong elevation of phosphon-ATM activity in response to high drug may trigger more extensive phosphorylation of Mdm2, thus promoting its auto-degradation and restraining Mdm2 expression close to the basal level. To determine whether Mdm2 degradation is enhanced at high etoposide concentration and whether this degradation process is ATM dependent, we measured the half-life of Mdm2 protein in U-2 OS cells by western blotting, using cycloheximide to inhibit overall protein translation. As shown in Figure 3a, levels of Mdm2 decreased with a half-life of about 1.96 hours in control U-2 OS cells (no etoposide treatment), 2.02 hours under 1μM etoposide (i.e. low drug) and 0.31 hours under 100μM etoposide (i.e. high drug). That is, the rate of Mdm2 degradation increased by about 6.5 fold at high drug as compared to that of low drug and control. Moreover, combinatorial treatment of U-2 OS cells with 100μM etoposide and 20μM KU55933, a small-molecule inhibitor of ATM, significantly prolonged the half life of Mdm2 to 1.16 hours, indicating that the accelerated Mdm2 degradation in U-2 OS cells at high etoposide concentration is mainly mediated by increased ATM activity (Fig. 3b). Overall, our data suggest that enhanced Mdm2 degradation mediated by increased ATM activity under high drug is the key mechanism underlying the attenuated Mdm2 up-regulation. The combined inhibitory interaction of ATM-Mdm2 and Mdm2-p53 thus form a double-negative, i.e., positive, feed-forward control that renders strong, monotonically increasing p53 in U-2 OS cells in response to high drug.

**Figure 3.**
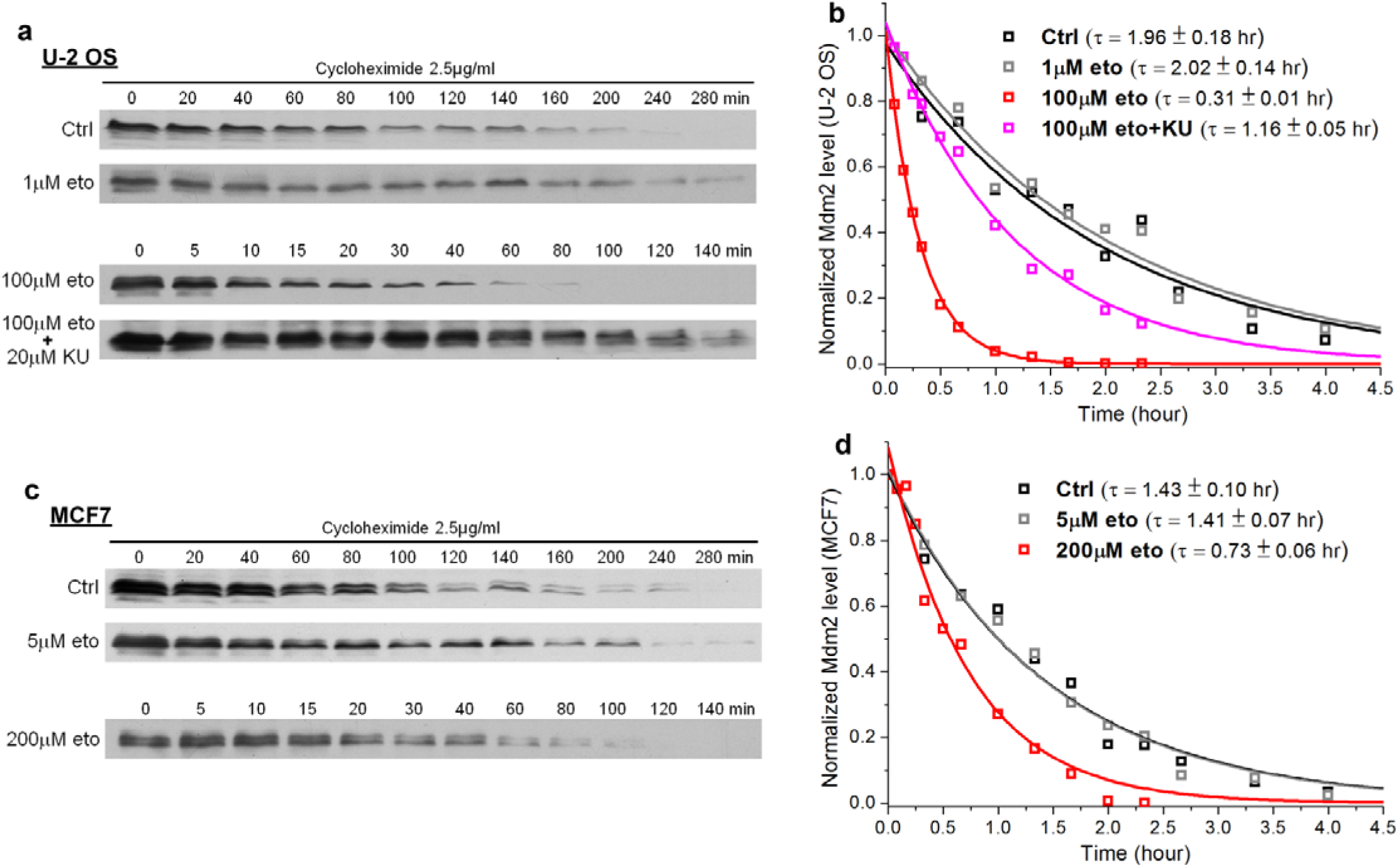
Dose-dependent Mdm2 expression is mainly regulated by ATM-mediated Mdm2 degradation. (a) Mdm2 degradation kinetics in U-2 OS cells at the indicated treatment condition plus cycloheximide (2.5μg/ml), were measured by western blot analysis of Mdm2 level at 12 selected time points (unit: minute). Actin, which served as loading control, was not shown, as loading were similar in all samples. b) Exponential fit of the Mdm2 degradation kinetics averaged from three independent sets of western blots and the derived Mdm2 half life τ at the distinct treatment condition. (c) Mdm2 degradation kinetics and (d) their respective exponential fits in MCF7 cells at the indicated treatment condition plus cycloheximide (2.5μg/ml). Data were averaged from three independent sets of western blots.

To confirm the inhibitory ATM-Mdm2 interaction was not activated to a substantial level to suppress Mdm2 expression in MCF7 under high drug, we also measured the half-life of Mdm2 in MCF7 cells. The Mdm2 half-life reduced by less than 2 fold in MCF7, as etoposide concentration increased, i.e., 1.43 hours in untreated cells, 1.41 hours under 5μM etoposide and 0.73 hour under 200μM etoposide (Fig. 3b), indicating the auto-degradation of Mdm2 promoted by ATM phosphorylation is significantly less in MCF7 than U-2 OS. This can be attributed to the significantly lower level of phospho-ATM (a surrogate of ATM activity) in MCF7, resulting from the low expression of total ATM.

### Mathematical modeling of p53 dynamics

In order to quantitatively understand how the feed-forward loop of ATM-Mdm2-p53 together with the negative feedback loop of Wip1-p53 collectively controls dose-dependent bimodal p53 dynamics, and to investigate cell type differences, we performed a computational analysis. We established a four-component model, consisting of ATM, p53, Mdm2 and Wip1 (Fig. 4a and 4b). We implemented the standard negative feedback loop between p53 and Mdm2, both of which can be phosphorylated by ATM and dephosphorylated by Wip1. As Wip1 is also a transcriptional target of p53, a second negative feedback loop was implemented between p53 and Wip1. Phosphorylation of p53 changes its stability as well as transcription activity. And phosphorylation of Mdm2 by ATM attenuates its inhibitory interaction with p53 as well as promotes its auto-degradation, thus forming a coherent feed-forward loop between ATM, Mdm2 and p53. Dynamics of this model are governed by the following delay differential equations (DDEs):
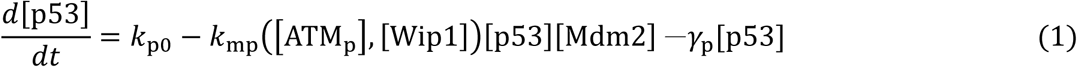

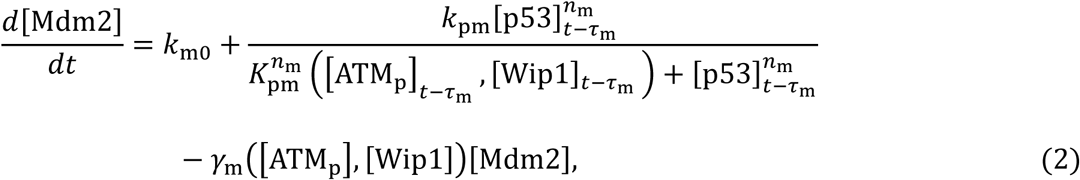

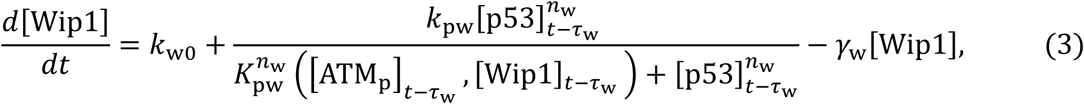

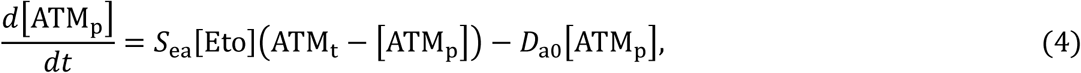

**Figure 4.**
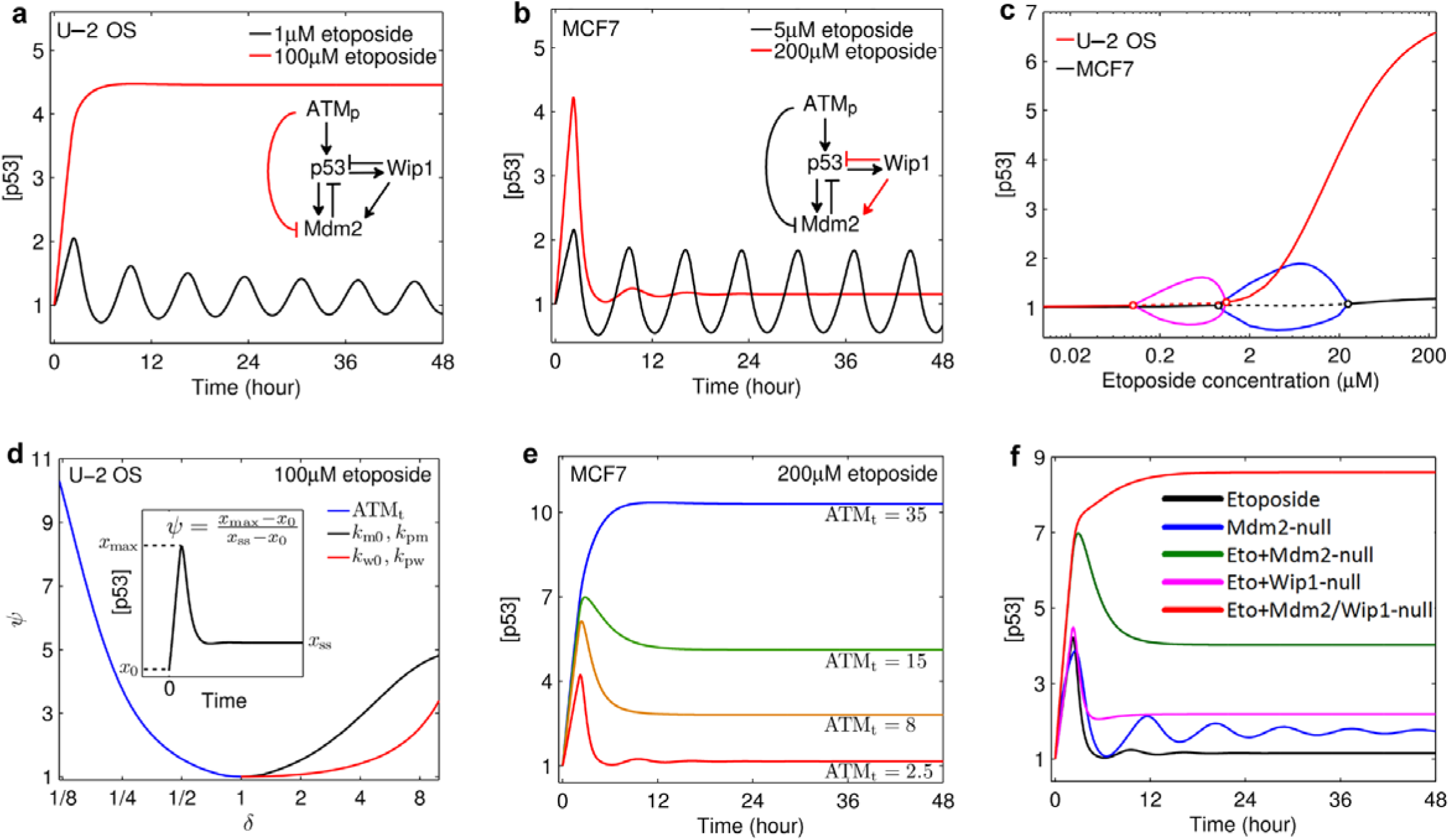
Bimodal p53 dynamics are regulated by a four-component regulatory module that consists of ATM, p53, Mdm2 and Wip1. (a, b) Simulated single-cell dynamics of p53 and the corresponding four-component module. The black and red trajectories are p53 level as a function of time under 1μM (5μM) and 100μM (200μM) etoposide in (a) U-2 OS and (b) MCF7 cells, respectively. (c) Bifurcation diagram of p53 concentration versus etoposide concentration. The red and black curves denote the respective steady states for U-2 OS and MCF7 cells. The solid and dotted curves separately denote the stable and unstable states. The circles denote the Hopf bifurcation points. The magenta (blue) curves denote the maxima and minima of [p53] in the limit cycles for U2OS (MCF7) cells. X-axis is on log scale. (d) Dependence of p53 dynamics on ATM_t_, *k*_m0_, *k*_pm_, *k*_w0_ and *k*_pw_. *k*_pm_ and *k*_w0_ are changed with the same *δ* simultaneously, so as *k*_w0_ and *k*_pw._ (e) Effects of total amount of ATM on p53 dynamics in MCF7 under 200μM etoposide. (f) Simulated p53 dynamics in MCF7 cells under the following conditions: 200μM etoposide (black), Mdm2 inhibition (blue), 200μM etoposide + Wip1 inhibition (magenta), 200μM etoposide + Mdm2 inhibition (green), and 200μM etoposide + inhibition of Mdm2 plus Wip1 (red).

where [] denotes dimensionless concentrations of the total proteins (p53, Mdm2, Wip1) or active form of the protein (phospho-ATM, denoted by ATM_p_) (refer to supplementary information for derivation of the equations). These four equations are derived using the rapid equilibrium assumption to simplify the phosphorylation and dephosphorylation of p53 and Mdm2. *k*_p0_, *k*_m0_, and *k*_w0_ are the basal production rate of p53, Mdm2, and Wip1, respectively. The p53-induced production rates of Mdm2 and Wip1 are characterized by Hill function, where *τ*_m_ and *τ*_w_ are the respective time delays in Mdm2 and Wip1 production. Moreover, we assume a linear basal degradation of p53, Mdm2 and Wip1, while Mdm2-mediated degradation of p53 is described by a second-order reaction. For simplicity, only phosphorylated and unphosphorylated monomers of ATM (ATM_p_ and ATM_u_) are included in the model. The total concentration of ATM, [ATM_t_] = [ATM_u_] + [ATM_p_], is considered a constant. Given that ATM and Wip1 can change the stability of p53 and Mdm2 as well as the transcription activity of p53, *k*_mp_, *K*_pm_, *K*_pw_ and *γ*_m_ are taken as functions of [ATM] and [Wip1] (refer to supplementary information for expression of these coefficients). To simulate the ODEs for the two different cell lines, we selected parameter values based on the western blot results (Supplementary Table S1 and S2). In particular, U-2 OS cells are set to have a higher level of total ATM, while MCF7 cells have higher level of Mdm2 and Wip1.

Simulation of the four-component model at low and high etoposide concentration produced the same cell line-specific bimodal p53 dynamics as observed in the single cell imaging experiments (Fig. 4a and 4b). p53 oscillation is activated in both U-2 OS and MCF7 cells at low drug, where steady states lose stability and limit cycles appear in the bifurcation diagram (Fig. 4c). Under high etoposide concentration, the steady-state p53 in U-2 OS cells can reach a high level because the high level of ATM_p_ strongly phosphorylates Mdm2 and promotes its degradation. Mdm2 thus remains largely at the basal level, allowing p53 to accumulate to a high level (Fig. S1). In contrast, the steady state of p53 in MCF7 only rises slightly at high drug (Fig. 4c). Hence, p53 tends to return to a low level after transient elevation, resulting in the dynamic mode of a single pulse. This dynamic mode can be explained by low ATM_p_ activity due to the low level of total ATM in MCF7, which is insufficient to enhance Mdm2 degradation significantly. As Mdm2 degradation does not outweigh its production, Mdm2 increases to a high level after a rapid degradation, which drives p53 back to a low level and keeps it at that low level (Fig. S1).

Using this model, we were able to computationally investigate the quantitative dependence of p53 dynamics on ATM, Mdm2 and Wip1 in response to high etoposide concentration, where U-2 OS and MCF7 differ. To distinguish dynamics of monotonic increase and single pulse, we first defined a quantity *ψ* as follows to quantify the degree, to which p53 dynamics resemble a typical single pulse,
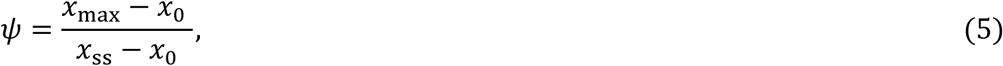

As shown in the inset of Figure 4d, *x*_0_ is the initial level of p53 (i.e., steady-state value under no drug treatment), *x*_max_ is the maximal p53 level induced by etoposide, and *x*_ss_ is the steady-state level of p53 under drug. Note that *ψ* = 1 indicates that p53 dynamics are monotonic increase and *ψ* > 1 indicates a single pulse. A larger *ψ* corresponds to p53 dynamics with features closer to a typical single pulse. We first simulated p53 dynamics for U-2 OS cell under perturbations of total ATM (ATM_t_) and the production rates of Mdm2 (*k*_m0_ and *k*_pm_) and Wip1 (*k*_w0_ and *k*_pw_), by increasing or decreasing their original values by *δ* folds. Notably, reducing total ATM or increasing the transcription rates of Mdm2 and Wip1 can change p53 dynamics in U-2 OS from monotonic increase to a single pulse, indicating these are viable routes to evolve drug resistance. Moreover, decrease of total ATM exhibits a greater effect on altering p53 dynamic than increasing the production rates of Mdm2 and Wip1, further validating that ATM expression, which determines the activation level of the positive feed-forward of ATM-Mdm2-p53, is the most critical factor that controls cell-type variation in p53 dynamics.

Next we simulated how p53 dynamics can be altered from single pulse to monotonic induction in MCF7 cells (Fig. 4e and 4f). Considering monotonic p53 induction activates extensive etoposide-induced cell death, changing single-pulse p53 to monotonic increase may be an effective way to enhance the cell death response of MCF7. As expected, increasing the level of total ATM in the model parameter can change p53 dynamics from a single pulse to monotonic increase in MCF7 (Fig. 4e). Inhibition of Mdm2 activity from the model has a notable effect on prolonging the duration of p53 pulse and increasing the steady-state level of p53 (i.e., a higher plateau that the pulse falls back on) at high drug, but p53 dynamics remain a single pulse (Fig. 4f). Removal of Wip1 does not significantly affect p53 dynamics. It is only under the simultaneous inhibition of Mdm2 and Wip1 that p53 dynamics change from a single pulse to monotonic induction in MCF7 under high drug. These results indicate both Mdm2 and Wip1 play important inhibitory role on p53 dynamics, and the inhibitory strength exerted by the negative feedback of Mdm2-p53 is stronger than that by Wip1-p53. In other words, Mdm2-p53 functions as the primary inhibitory feedback, while Wip1-p53 serves as a secondary inhibitory control that is critical when the primary negative feedback of Mdm2-p53 is inhibited or attenuated.

### Sensitization of MCF7 cells to drug-induced cell death by modulation of p53 dynamics

To validate the modeling results regarding the control of p53 dynamics and examine whether change of p53 dynamics can enhance cellular response to etoposide, in particular the cell death response in MCF7 at high drug, we performed single cell imaging experiments of MCF7 treated with 200μM etoposide plus inhibition of Mdm2, Wip1 or both. We used Nutlin-3, a small-molecule inhibitor of Mdm2, to inhibit Mdm2 activity. Wip1 was removed by RNAi knockdown. Knockdown efficiency was generally more than 90% and Wip1 knockdown alone induced little cell death (< 5%) up to 72 hours of experiment. As shown in Figure 5a, loss of Wip1 did not alter the single-pulse dynamics of p53 induction. Upon Mdm2 inhibition by Nutlin-3, p53 dynamics changed from a short single pulse (duration of 8-12 hour) to a long pulse (duration of 18-25 hours), although the steady-state level of p53 was lower than that predicted by the model simulation. Strong, monotonic induction of p53 was observed in MCF7 under the combinatorial treatment of Nutlin-3, Wip1 knockdown and 200μM etoposide, confirming the simulation results. Moreover, we found change of p53 dynamics to a higher induction mode significantly enhanced and accelerated drug-induced cell death in MCF7 cells (Fig. 5b). The monotonically increasing p53 resulted from the triple treatment triggered more than 80% cell death in MCF7 after 36 hours, as compared to less than 10% death in etoposide treatment alone for the same time duration, illustrating that altering p53 dynamics is sufficient to sensitize MCF7 cells to etoposide-induced cell death and that Mdm2 and Wip1 are viable combinatorial targets to enhance response of resistant cancers to DNA damaging drug.

**Figure 5.**
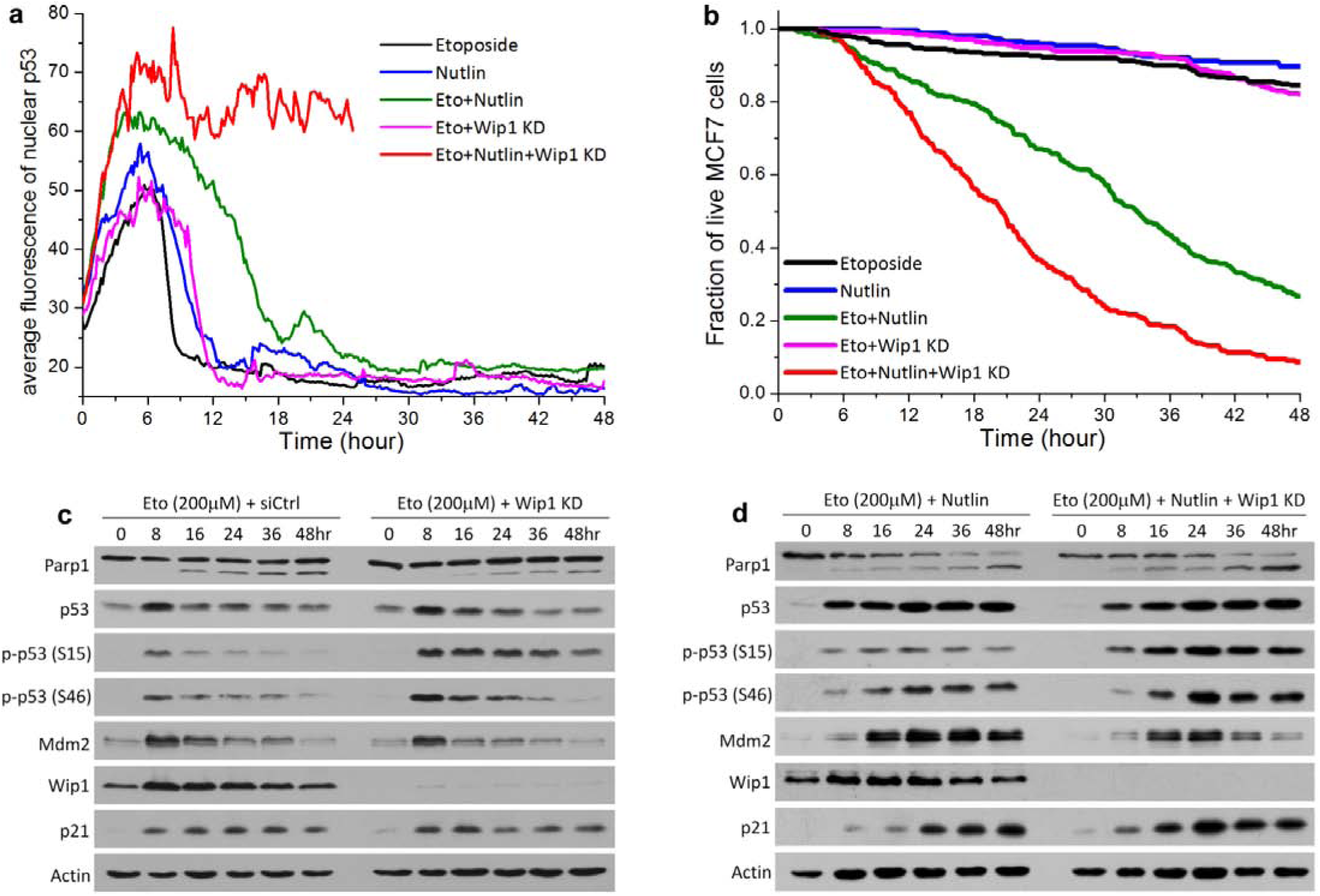
Inhibition of Mdm2 and Mdm2 plus Wip1 alters p53 dynamics and subsequent drug-induced cell death. (a) Representative single-cell trajectories of p53-Venus fluorescence in MCF7 cells under the indicated treatment, including etoposide alone (200μM), Nutlin-3 (10μM) alone, etoposide + Nutlin-3, etoposide + Wip1 knockdown (kd) and etoposide + Nutlin-3 + Wip1 kd. The abrupt end of p53 trajectory under etoposide + Nutlin-3 + Wip1 kd corresponds to the time of death. (b) Cumulative survival curves of MCF7 cells under the indicated treatments. Data were averaged from three independent imaging experiments and the total number of cells analyzed ranges from 62 to 132, varied between conditions and experiments. Cell death was scored morphologically based on time-lapse movies, and kinetics of cell death was plotted as cumulative survival curves. (c, d) Western blot comparison of selected proteins/protein modifications of MCF7 cells treated with (c) 200μM etoposide plus control siRNA vs. Wip1 knockdown, or (d) 200μM etoposide + 10μM Nutlin-3 vs. etoposide + Nutlin-3 + Wip1 kd.

Further western blot analysis confirmed not only the single cell imaging results regarding differential p53 dynamics and cell death response in MCF7 cells but also that higher p53 induction level corresponded to higher p53 activity, e.g., in up-regulating p21 (Fig. 5c and 5d). Moreover, it showed that Wip1 knockdown strongly enhanced p53 phosphorylation, e.g., at Serine 15 and Serine 46. Phosphorylation of p53 at Serine 15 is known to stabilize p53 [10, 27], and phosphorylation at Serine 46 particularly activates the transcription activity of p53 to up-regulate pro-apoptotic genes [28-30]. Therefore, the negative feedback of Wip1-p53 likely attenuates p53 induction level by destabilizing p53 and further decreases cell death response to etoposide by attenuating p53 transcriptional activation of pro-apoptotic genes.

## Discussion

Our study not only elucidates a dynamic mechanism underlying variable sensitivity to DNA damaging drug but also provides new insight to understand cell-type variation in p53 pathway-mediated DNA damage response. Variation in cell death response to cytotoxic chemotherapy is possibly the most crucial factor that distinguishes sensitive and resistant tumors, as clinical response of tumor regression or delay in tumor growth often correlates with the cell death response [31, 32]. Our data showed that the dynamic mode of p53 induction, especially at regime of high concentration of DNA damaging drug (i.e., high level of DNA damage), determines the sensitivity of p53-wild-type cancer cells to drug-induced cell death. Resistance to the cell death response arises from mechanism not due to obvious genetic mutation, but through attenuating the output of a collective regulatory module that controls p53 dynamics. In MCF7 cells, the modular output is damped mainly by down-regulating ATM. We suspect perturbations of other modular features (e.g., enhancement of Mdm2 and Wip1 transcription rates) or additional regulatory components beyond this core module, which promote pulsing p53 upon drug treatment, may contribute to drug resistance of other tumor types. Our results also suggest combinatorial targeting of Mdm2 and Wip1 is a promising strategy to combat this type of drug resistance.

The fact that we found down-regulation of ATM reduced sensitivity to DNA damaging drug puts a cautious note on combining ATM inhibitor with DNA damaging chemotherapeutics for cancer treatment [33]. Loss of ATM activity was first found to sensitize cells to ionizing radiation (IR), sparking the interest to develop ATM inhibitor as anticancer drug, particularly as combinatorial therapy with IR and DNA damaging drugs [34]. Early data showed tumor cells with defective p53 was more pronouncedly sensitized by ATM inhibitor to IR and chemo-drugs [35, 36], while a recent study concluded the sensitizing effect was significant in both p53-wild-type and p53-defficient cells [37]. It remains unclear whether and how the functional status of p53 affects response to ATM inhibitor. Our study showed attenuation of ATM activity desensitizes, rather than sensitizes, the response of p53-wild-type cancer cells to DNA damaging drug. Although down-regulation of ATM activity may not give rise to exactly the same phenotypic response as that of complete inhibition of the kinase, our results do argue against combining ATM inhibitors with chemo-drugs for tumors with wild-type p53.

By exploring a large range of DNA damage dose, we characterized p53 dynamics and drug response at a dynamical regime that is more associated with cell death. The typical etoposide dose used in patients is 100mg/m^2^/day, which results in a peak plasma drug concentration of about 20μg/ml, i.e., 34μM [38]. As shown in Figure 1d, this dose level of etoposide is sufficient to activate monotonic p53 induction and cell death in 60-80% of U-2 OS cells, suggesting the dose response of U-2 OS is possibly informative of sensitive tumors in the clinical context. At the same dose level, less than 5% cell death was induced in MCF7, validating it as a model for resistant cancer. The resistance mechanism that we elucidated for MCF7 may help to guide identification of some resistant tumors. Less clear is how normal cells respond to etoposide at this treatment dosage. In order for the cytotoxic therapy to work, a favorable efficacy-to-toxicity ratio (‘therapeutic index’) has to be achieved, i.e., the drug needs to kill tumor without killing the patient. Normal cells obviously have wild-type p53, and our results suggest that retraining p53 dynamics to the pulsing mode protects cells from drug-induced cell death. We do not yet know how the regulatory module that controls p53 dynamics differs between normal and cancer cells and whether it is possible to develop combinatorial target that can achieve a better therapeutic index, e.g., by protecting normal cells or sensitizing cancer to respond to lower drug dose. Nonetheless, our results point to modulation of p53 dynamics as a new possible angle to explore new drug target and drug combination.

On the basic biology of signaling molecule dynamics and pathway control, our data provide new evidence that a major function of p53 pulsing, both in the form of periodic pulsing and single pulse, is to suppress p53 induction level and its activity to trigger cell death. At time of low DNA damage, p53 oscillation retrains p53 induction and its pro-apoptotic activity at low level, which allows damage repair and cell survival. Upon high DNA damage, single-pulse p53 prevents p53 induction to a high level that can activate rapid and extensive cell death, again enabling cell survival despite massive DNA damage. Periodic pulsing of p53 dynamics is extensively characterized before, with most studies attributing it to a time-delay negative feedback loop between p53 and Mdm2 [39, 40]. The dynamic mode of single-pulse p53 was also previously analyzed for MCF7 cells in response to transient UV irradiation and was attributed to differential activation of the phosphatase, Wip1 [18]. However, our results showed Wip1 did not play a significant role in engendering single-pulse p53 in the model system that we studied. Both experimental and computational analysis of the ATM/p53/Mdm2/Wip1 regulatory module showed attenuation of the positive feed-forward loop of ATM-Mdm2-p53 is the key to generate suppressive single-pulse p53 dynamics at high DNA damage. It appears mechanisms that activate even the same p53 dynamic mode could differ depending on the stimulus. To unravel how the regulatory module that controls p53 dynamics may be differentially activated in a stimulus-dependent manner, we will need to investigate and compare single-cell p53 dynamics in response to a wider variety of stimuli, such as ribosomal stress and hypoxia, in follow-up studies.

## Materials and Methods

### Cell culture

Cell lines were purchased from American Type Culture Collection (ATCC, USA) and cultured under 37°C and 5% CO_2_ in appropriate medium supplemented with 10% Fetal Calf Serum (FCS), 100U/ml penicillin and 100μg/ml streptomycin. U-2 OS was maintained in McCoy’s; and MCF7 was maintained in RPMI. To generate fluorescent reporter cells for live-cell imaging of real-time p53 dynamics, we infected U-2 OS and MCF7 cells with lenti-viruses encoding an established p53-Venus reporter and selected isogenic clones. We selected isogenic clones that exhibited dose response most similar to their respective parental line for conducting the study. The p53-venus reporter construct, consisting of wild-type p53 fused to a yellow fluorescent protein, Venus, was a generous gift from Dr. Galit Lahav at the Department of Systems Biology, Harvard Medical School.

### Chemicals and reagent

Etoposide, Nutlin-3 and the ATM kinase inhibitor, KU55933, were purchased from Tocris. siRNA for knocking down Wip1 (UUG GCC UUG UGC CUA CUA A) was custom synthesized by Dharmacon. Dharmacon On-Target plus siControl (#D-001810-01) was used as non-targeting siRNA control. Both siRNAs were used at final concentration of 40 nM and siRNA transfections were performed using Lipofectamine (Thermo Fisher) according to manufacturers’ instructions. Experiments were conducted after 48 hrs of gene silencing.

### Time-lapse microscopy

Cells were plated in 35 mm imaging dish (μ-dish, ibidi, Germany) and cultured in phenol red-free CO_2_-independent medium (Invitrogen) supplemented with 10% FCS, 100U/ml penicillin and 100μg/ml streptomycin. Cell images were acquired with the Nikon TE2000-PFS inverted microscope enclosed in a humidified chamber maintained at 37°C. Cells were imaged every 10 minutes using a motorized stage and a 20X objective (NA=0.95).

For data plotted in the phenotype distributions, we viewed and analyzed the images manually, using the MetaMorph software (Molecular Dynamics). From phase-contrast images, we scored by morphological tracking: interphase (by flat morphology), entry into mitosis (by cell rounding), cell division (by re-spreading and splitting), cell-cycle arrest (by absence of cell division for 72 hours) and cell death (by blebbing followed by cell lysis). The dynamic mode of nuclear p53 was scored based on the p53-Venus fluorescence in the nucleus. For quantifying the single-cell p53 traces, we used automatic cell tracking program that we developed using Matlab. The program consists of image analysis procedures that sequentially segment the individual cells, track them in time, identify the nucleus and measure the p53 fluorescence intensity in the nucleus.

### Western Blot Analysis

Cell lysates were obtained using LDS sample buffer (NuPAGE, Invitrogen). Proteins were resolved on 10-15% Tris-glycine gels and transferred onto PVDF membranes. Blots were probed with commercial primary antibodies and chemiluminescent detection using ECL-Prime (Amersham). Antibodies: PARP1 (#9542), phospho-p53 (S15) (#9284), ATM (#2873), p21 (#2947) and Puma (#4976), were purchased from Cell Signaling; p53 (#sc-126), Mdm2 (#sc-965) and Wip1 (#sc-20712) from Santa Cruz; γH2A.X (#06-570) from Millipore; phospho-ATM (S1981) (#ab81292) and phospho-p53 (S46) (#ab76242) from Abcam. Anti-actin (#A5316) from Sigma was used as a loading control.

### Mathematical model and computational analysis

Detailed description of the four-component model for the core p53 regulatory module and computational analysis of this module in terms of differential activation of bimodal p53 dynamics in response to variable DNA damage signal can be found in the Results section and supplementary information.

## Author contributions

J.S. and F.L. designed the study; R.Y., Y.Z. and Y.L. performed experiments; B.H. performed the computational analysis; R.Y., B.H. and J.S. analyzed the data; R.Y., B.H., F.L. and J.S. wrote the paper.

## Acknowledgements

We thank Dr. Galit Lahav (Department of Systems Biology, Harvard Medical School) for the p53-Venus lenti-viral vector. This work was supported by the Hong Kong Research Grant Council (#N_HKBU215/13 and #T12-710/16-R) to J. Shi and National Science Foundation of China (#31361163003) to F. Liu. The authors declare no conflict of interest.

